# High performance liquid chromatography-based metabolomics of *Psidium guajava* Linn. leaf extracts

**DOI:** 10.1101/2022.01.25.477015

**Authors:** Manasi S. Gholkar, Poonam G. Daswani, Vidhya V. Iyer, Tannaz J. Birdi

## Abstract

**Background:** Over the years, a number of methods have been introduced in the field of herbal medicine. Amongst these, metabolomics has rapidly emerged as a method of choice due to its wide applicability and versatility. High performance liquid chromatography (HPLC) is a preferred method for fingerprinting with advantages such as easy and wide availability, and relatively low maintenance cost. The current study used HPLC profiling and attempted to correlate it to anti-diarrhoeal activity of *Psidium guajava* (guava) extracts.

**Methods:** Ninety samples of guava leaves were collected from three locations in Maharashtra, India over three seasons. Hydroalcoholic extracts (50:50, ethanol:water) were prepared and the HPLC chromatogram of all 90 extracts obtained. A total of eight bioassays representing the important features of diarrhoeal pathogenesis were performed and the extracts were differentiated as ‘good’ or ‘poor’ according to cut-offs for each assay. The numerical data of the bioassays and the HPLC chromatogram was correlated using suitable mathematical models comprising Principal component analysis (PCA), orthogonal partial least squares (OPLS, regression) and orthogonal partial least squares discriminant analysis (OPLS-DA).

**Results:** OPLS-DA showed good seasonal and regional segregation of extracts. For most of the bioassays, PCA was unsuccessful in showing significant discrimination. Hence, OPLS plots were further developed; differentiation of good and poor extracts among the 90 extracts was successfully demonstrated for antibacterial activity against *Shigella flexneri* and *Vibrio cholerae*, inhibition of invasion of enteropathogenic *Escherichia coli* into HEp-2 epithelial cells, and cholera toxin (CT) production by *V. cholerae*. However, for other assays, namely, inhibition of adherence of *E. coli*, invasion of *S. flexneri* into HEp-2 cells, and inhibition of *E. coli* labile toxin (LT), subsets of 90 extracts were selected to demonstrate a significant correlation.

**Conclusion:** HPLC-based metabolomics has the potential to differentiate good and poor activities of guava leaf extracts. This approach can be extended for identifying phytoconstituents responsible for the anti-diarrhoeal activity of guava leaf and help standardization of crude extracts.

## Introduction

Plants synthesize chemically diverse molecules; this coupled with complexity of the constituents make phytochemical profiling and estimation of active compounds challenging as well as time consuming (Sumner et al., 2003). Over the years, a number of analytical methods have been developed for detailed profiling of herbal drugs. Chromatographic techniques such as thin layer chromatography (TLC) and high performance liquid chromatography (HPLC) are the most commonly used methods for obtaining chemical fingerprints and identification of phytoconstituents in plant extracts. Use of more sophisticated spectroscopic techniques such as nuclear magnetic resonance (NMR), infra-red (IR) spectroscopy have also gained popularity (Marston, 2007).

Chromatography which was initially employed as a simple tool for pigment separation has evolved over the years as a versatile technique capable of dealing with complex analytical and purification problems in phytochemical research (Marston 2007). Amongst the various chromatographic techniques, the area of HPLC has seen remarkable advances and is now considered as a powerful tool in analytical chemistry. HPLC has several advantages in the analysis of herbal medicines such as high sensitivity, good resolution and linearity, ability to analyze multiple constituents and ease of automation making it suitable for obtaining a fingerprint (Zhong et al, 2009). It has the scope of separating, identifying and quantitating compounds present in a sample. HPLC can be used for the qualitative as well as quantitative detection of the active or marker compounds. For example HPLC has been used for detection of alkylamides from *Echinacea species* that contains a number of isomeric and structurally similar alkylamides and for obtaining fingerprints of *Panax notoginseng* obtained from different sources (Bruni et al., 2018; Xu et al., 2019).

Leaves of *Psidium guajava* (guava) Linn. are globally used as a traditional medicine to treat diarrhoea and other gastro-intestinal ailments, including dysentery and amebiasis (Daswani et al., 2017, Kafle et al., 2018). Extensive work undertaken at Foundation for Medical Research, including *in vitro* and *in vivo* studies, followed by a clinical trial in adults with uncomplicated diarrhoea, established guava leaves as a promising anti-diarrhoeal remedy having a wide spectrum of activity (Birdi et al., 2010, 2011, 2014, 2020; Brijesh et al., 2011; Gupta and Birdi, 2015). The guava leaves are rich in phytoconstituents and a number of constituents including carotenoids, triterpenoids and flavonoids, sesquiterpenes, saponins, sterols, phenolics, coumarins have been reported by several researchers (Anand et al. 2016; Camarena-Tello et al. 2018; Díaz-de-Cerio et al. 2015; Metwally et al. 2010). Amongst the various constituents, phenolics such as protocatechuic acid, caffeic acid, gallic acid, ferulic acid and quercetin are abundantly present in the leaves (Koreim et al., 2019; Hsieh et al., 2007; Chen et al., 2007).

The promising anti-diarrhoeal activity of guava leaves and lack of detailed information on its functional compounds made this plant a good choice in our studies for the development of a prototype for standardization of crude plant extracts. Standardization of crude extracts may be attempted by two main approaches. The first would be obtaining a representative fingerprint which is considered distinctive and forms a benchmark for a particular extract especially when the identity of the active principle(s) is unknown (Rajani et al., 2008). The second approach involves identification of marker compounds which necessarily do not always correlate with biological activity.

The utility of a spectral fingerprint toward standardization of crude extracts was successfully demonstrated in our previous study (Gholkar et al., 2021). The study attempted to develop a prototype for standardization of extracts used ^1^H NMR metabolic profiling by correlating peaks to anti-diarrhoeal activity of guava leaf extracts.

In the present study we have explored the utility of HPLC based metabolomics as an approach towards standardization of guava extract by possible correlation with the biological efficacy of the extract. Towards this, guava leaves were collected in different seasons from different locations and their HPLC profiles were acquired through a PDA detector. Eight bioassays representing important stages in diarrhoeal pathogenesis were undertaken with each batch of leaves collected and the results were correlated with the HPLC profiles using mathematical models.

## Materials and Methods

### Plant material and preparation of extract

Ninety samples of *Psidium guajava* (guava) leaves of the *Sardar* variety were collected, ten trees each, from three regions in Maharashtra (Shirwal, W; Rahata, R and Dapoli, Da) over three seasons (B, May 2013; C, October 2013 and D, March 2014). The leaves were washed, shade-dried, powdered and a hydro-alcoholic extract (50:50) prepared. Following the evaporation of the alcohol from the extracts, the aqueous portion was lyophilized and the resulting powder stored at −80°C. Extracts were reconstituted in distilled water before being use.

### Cell culture

HEp-2, a human laryngeal epithelial cell line procured from the National Centre for Cell Sciences, Pune, India was maintained in Dulbecco’s Modified Eagle’s Medium (DMEM Gibco) supplemented with 10% FCS (Bio-west) at 37°C in a 5% CO_2_ atmosphere. The cells were passaged at least twice a week.

### Bacterial strains

The following five bacterial strains were used for the study

1. Enteropathogenic *Escherichia coli* (EPEC) strain B170, serotype 0111:NH,
2. Enterotoxigenic *E. coli* (ETEC) strains B831-2, serotype unknown (heat labile toxin producer) (both obtained from Center for Disease Control, Atlanta, USA),
3. Entero-invasive *E. coli* (EIEC) strain EI34, serotype 0136:H- (kindly provided by Dr. J. Nataro, Veterans Affairs Medical Centre, Maryland, USA);
4. *Vibrio cholerae* Ogawa, serotype 01 (kindly provided by Dr. S. Calderwood, Massachusetts General Hospital, Boston, USA) and
5. *Shigella flexneriM9OT*, serotype 5 (kindly provided by Dr. P. Sansonetti, Institut Pasteur, France)

#### Bioassays

The methods for all the bio-assays as described below are well established. Additionally, these have been validated earlier using suitable positive controls (Birdi et al., 2010).

##### Effect on antibacterial activity

The agar dilution method was used to assess antibacterial activity against the five bacterial strains (Cruickshank 1975). Bacterial strains were plated on Mueller Hinton agar (Himedia laboratories, India),without extract (control) and containing 1000 μg/mL,150 μg/mL and 600 μg/mL (wt/vol) of the reconstituted extract (as test) for *E. coli, S.flexneri* and *V. cholerae* respectively. Two independent experiments were carried out in triplicate. Results are expressed as percent viability; colony-forming units (cfu) from the control plates were taken as 100%.

##### Effect on bacterial colonization

###### Bacterial adherence

Adherence of EPEC strain B170 to HEp-2 cells was assayed as described previously (Cravioto et al., 1998). In brief, cells cultured on glass coverslips for 48 h were infected with a log phase culture (5 × 10^7^/mL) of bacteria in DMEM alone (control) or 20 μg/mL of the extract in DMEM (test). Following a 3 h incubation, the non-adherent bacteria were washed off, the coverslips fixed using methanol and stained with toluidine blue (0.1% w/v). HEp-2 cells showing the characteristic adherent EPEC microcolonies and/or >5 adherent bacteria were counted under a microscope. Two independent experiments were carried out and for each assay duplicate coverslips were set up for test and control. Results are expressed as percent adherence; HEp-2 cells counted from the control were taken as 100%.

###### Bacterial invasion

Invasion of EIEC strain EI34 and *S. flexneri* intoHEp-2 cells was carried out using a protocol reported earlier (Vesikari *et al*.1982). For this assay, HEp-2 cells were grown in a 96-well tissue culture plate and infected with a log phase culture (10^8^/mL) of the bacteria in DMEM alone (control) or in DMEM containing 20 μg/mL of the extract (test). Following a 2 h incubation, the extracellular bacteria were washed off and the cells incubated with DMEM containing gentamycin (100 μg/mL) for additional 2 h to kill extracellular bacteria. Thereafter, the medium containing gentamycin was washed off and the cells lysed with chilled distilled water. The counts of the intracellular bacteria thus released were obtained by plating the cell lysate on nutrient agar. Two independent experiments were carried out with triplicate wells being set up for the test and the control. Results are expressed as percent viability with cfu from the control wells being regarded as 100%.

##### Effect on bacterial enterotoxins

Heat-labile toxin (LT) produced by ETEC and extracellularly released cholera toxin (CT) were assayed by the ganglioside monosialic acid enzyme-linked immunosorbent assay (GM1-ELISA) by using a previous method (Svennerholm and Wilkund, 1983). Briefly, bacteria were grown for 18-20 h in medium without (control) and with the extract to produce the toxin. LT was extracted by treating the bacteria with polymyxin B sulphate and CT was obtained as the culture supernatant. ELISA was carried out using anti-cholera toxin and HRP labelled swine immunoglobulins as the primary and secondary antibodies respectively. The color end product was read at optical density (OD) of 492 nm. Data was expressed as percent toxin produced and OD of the control was taken as 100%. Two independent experiments were carried out in triplicate for test and control. Results are expressed as percent toxin produced with OD of the control taken as 100%.

To study the effects of the extract on LT/CT production, bacteria were grown for 18-20 h in medium with (test) and without (control) extract. 100 μg/mL and 75 μg/mL were the extract concentrations used for LT and CT, respectively. To study the effects of the extract on CT binding to GM1, CT toxin from untreated *V. cholerae* supernatant was taken and added to GM1 in the presence and absence of the extract. The concentration of the extract to check the effect of the extract on the binding of the released CT was 50 μg/mL extract.

##### Determination of concentrations and cut-off limits for all extracts

Pilot experiments were performed for each of the bioassays to arrive at a concentration of the extract showing a wide spectrum of activity. Following calculation of the mean value of two experiments for each assay, cut-off limits were empirically set to mark extracts with good (higher inhibition of the parameter) and bad/poor (lower inhibition of the parameter) activities. Values between these cut-off limits were considered to be intermediate. The extract concentrations and cut-off limits differed for each bioassay and have been depicted in Table 1.

**Table 1:**
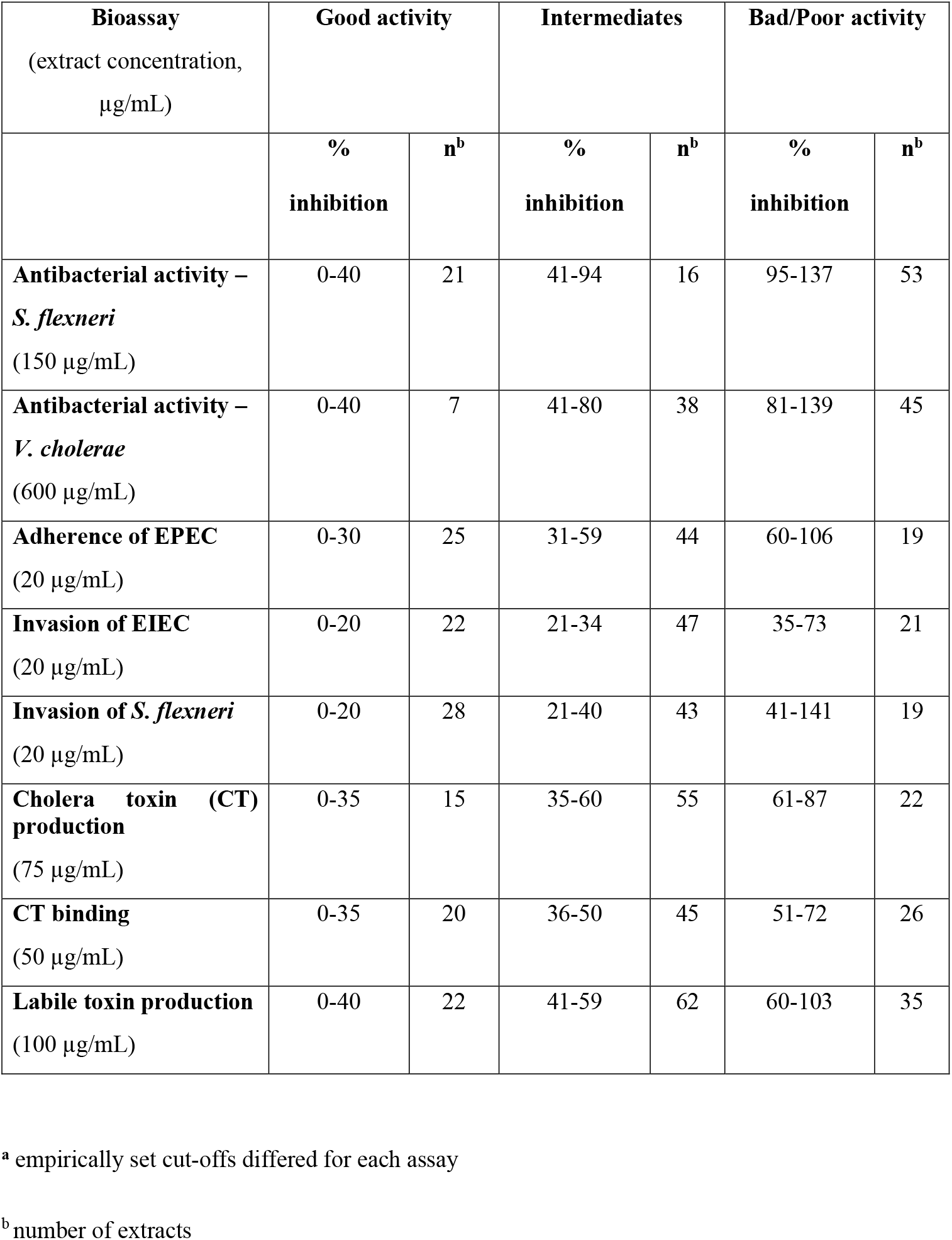
Activity-based classification of extracts based on the individual cut-offs^a^.

| **Bioassay**<br>(extract concentration,<br>μg/mL) | **Good activity** |  | **Intermediates** |  | **Bad/Poor activity** |  |
| --- | --- | --- | --- | --- | --- | --- |
|  | **%**<br>**inhibition** | **n<sup>b</sup>** | **%**<br>**inhibition** | **n<sup>b</sup>** | **%**<br>**inhibition** | **n<sup>b</sup>** |
| **Antibacterial activity –**<br>***S. flexneri***<br>(150 μg/mL) | 0-40 | 21 | 41-94 | 16 | 95-137 | 53 |
| **Antibacterial activity –**<br>***V. cholerae***<br>(600 μg/mL) | 0-40 | 7 | 41-80 | 38 | 81-139 | 45 |
| **Adherence of EPEC**<br>(20 μg/mL) | 0-30 | 25 | 31-59 | 44 | 60-106 | 19 |
| **Invasion of EIEC**<br>(20 μg/mL) | 0-20 | 22 | 21-34 | 47 | 35-73 | 21 |
| **Invasion of *S. flexneri***<br>(20 μg/mL) | 0-20 | 28 | 21-40 | 43 | 41-141 | 19 |
| **Cholera toxin (CT)**<br>**production**<br>(75 μg/mL) | 0-35 | 15 | 35-60 | 55 | 61-87 | 22 |
| **CT binding**<br>(50 μg/mL) | 0-35 | 20 | 36-50 | 45 | 51-72 | 26 |
| **Labile toxin production**<br>(100 μg/mL) | 0-40 | 22 | 41-59 | 62 | 60-103 | 35 |

<sup>a</sup> empirically set cut-offs differed for each assay
<sup>b</sup> number of extracts

### High-performance liquid chromatography (HPLC)

The extracts dissolved in acetonitrile at a concentration of 1 mg/mL, were chromatographed by gradient elution on a NEXERA XR LC-20AD system (Shimadzu, Japan) equipped with an ultraviolet-PDA detector (200-600 nm) and DGU-20A 5R degassing unit. A Poroshell 120 column was used (100 × 3 mm, 2.7 μ particle size) with a mobile phase of acetonitrile: acetic acid, which changed from 95:5 to 82:18 over 12 min, and then was maintained at 0:100 from 12 to 17 min, and then again changed to 95:5 until the end of the run at 20.01 min.

Spectra for all the 90 extracts were recorded using Photodiode-Array Detector (PDA) over wavelengths of 220 to 500 nm.

#### Statistical analysis

For all assays, percentages have been calculated by using the formula {(C or T)/C} x 100, where C is the mean value of the control group and T is the mean value for the test (extract) group. Results for all assays have been expressed as the mean ± standard deviation of the percentage values of all replicates from two independent experiments. Cut-off values determined for all assays were used to categorize all activities as good (G), poor (bad B) or intermediate (I).

The HPLC chromatograms converted to numerical form along with the values of the bio-assay were imported into SIMCA (14.0). The data were correlated using unsupervised Principal Component Analysis (PCA) using category labels (good G, poor or bad B, intermediate I). Additionally, supervised Orthogonal Partial Least Square-Discriminant Analysis (OPLS-DA) was carried out to observe the metabolic differences between the extracts with ‘good’ bioactivity and that with ‘poor’ bioactivity. OPLS-Regression Analysis (OPLS-RA) was favored over OPLS-DA models wherever significant results were not observed. In some cases, subsets of data had to be used to obtain significance. The plots generated in SIMCA were colored as per the activity under consideration and subjected to Pareto scaling.

## Results

### HPLC chromatograms

Spectra of all the hydroalcoholic extracts recorded using PDA detector over wavelengths of 220 to 500 nm were converted into a max plot. A representative max plot has been depicted in Figure 1.

**Fig. 1:**
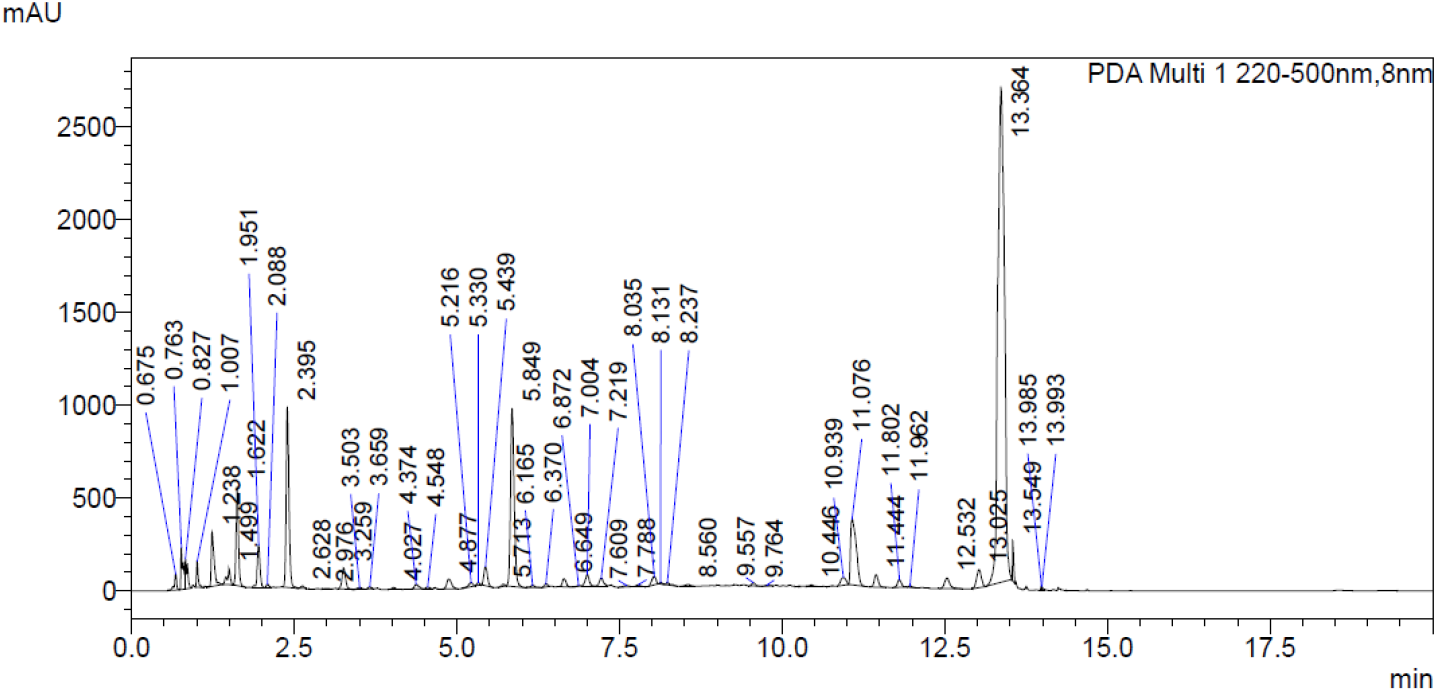
A representative MaxPlot of a *Psidium guajava* extract.

### Regional and seasonal differentiation

As seen in Figures 2 and 3, OPLS-DA showed clear differentiation between the extracts collected from different regions and seasons.

**Fig. 2:**
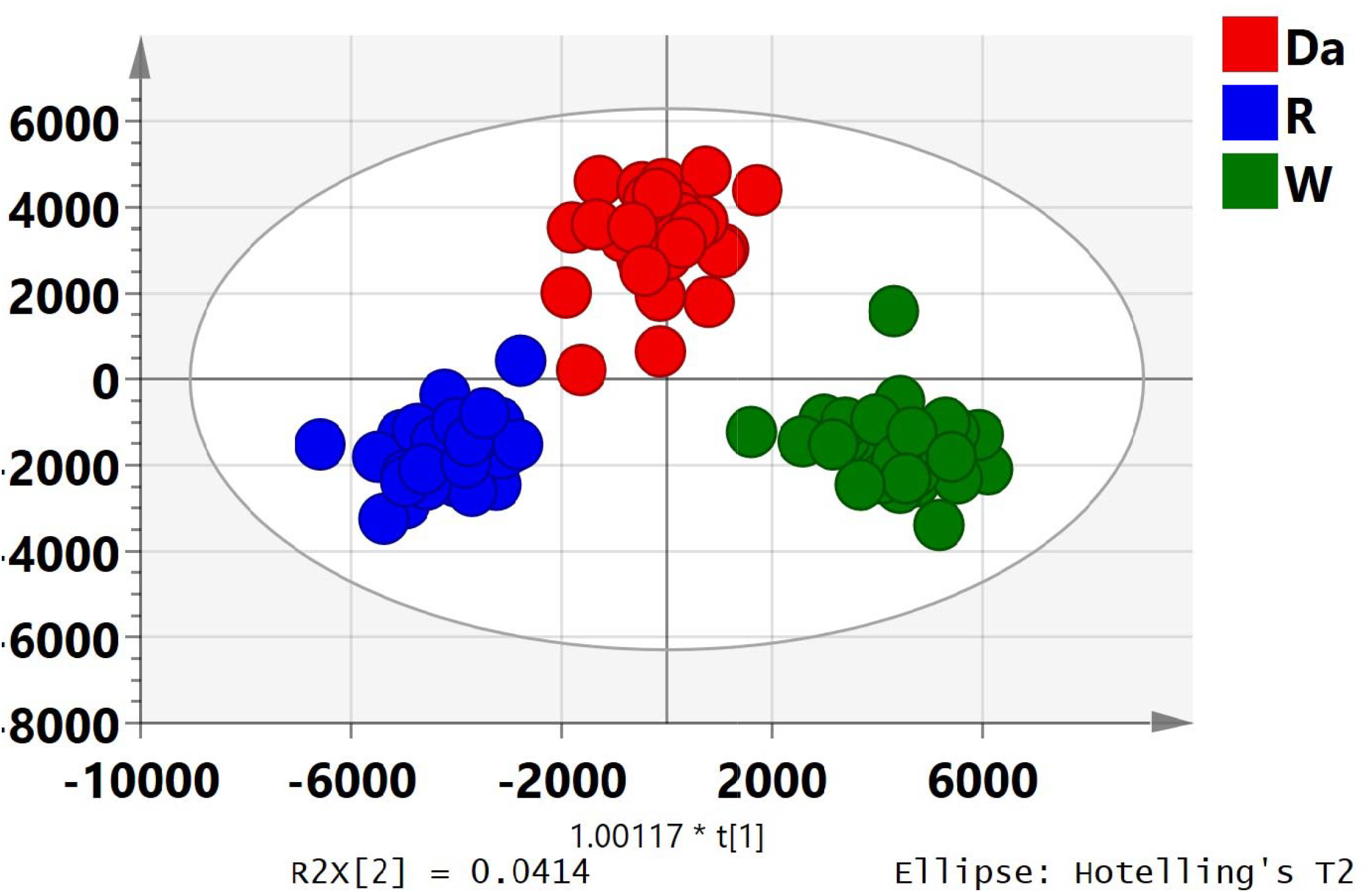
OPLS-DA plot for regional differentiation Region Da: Dapoli, Region R: Rahata, Region W: Shirwal

**Fig. 3:**
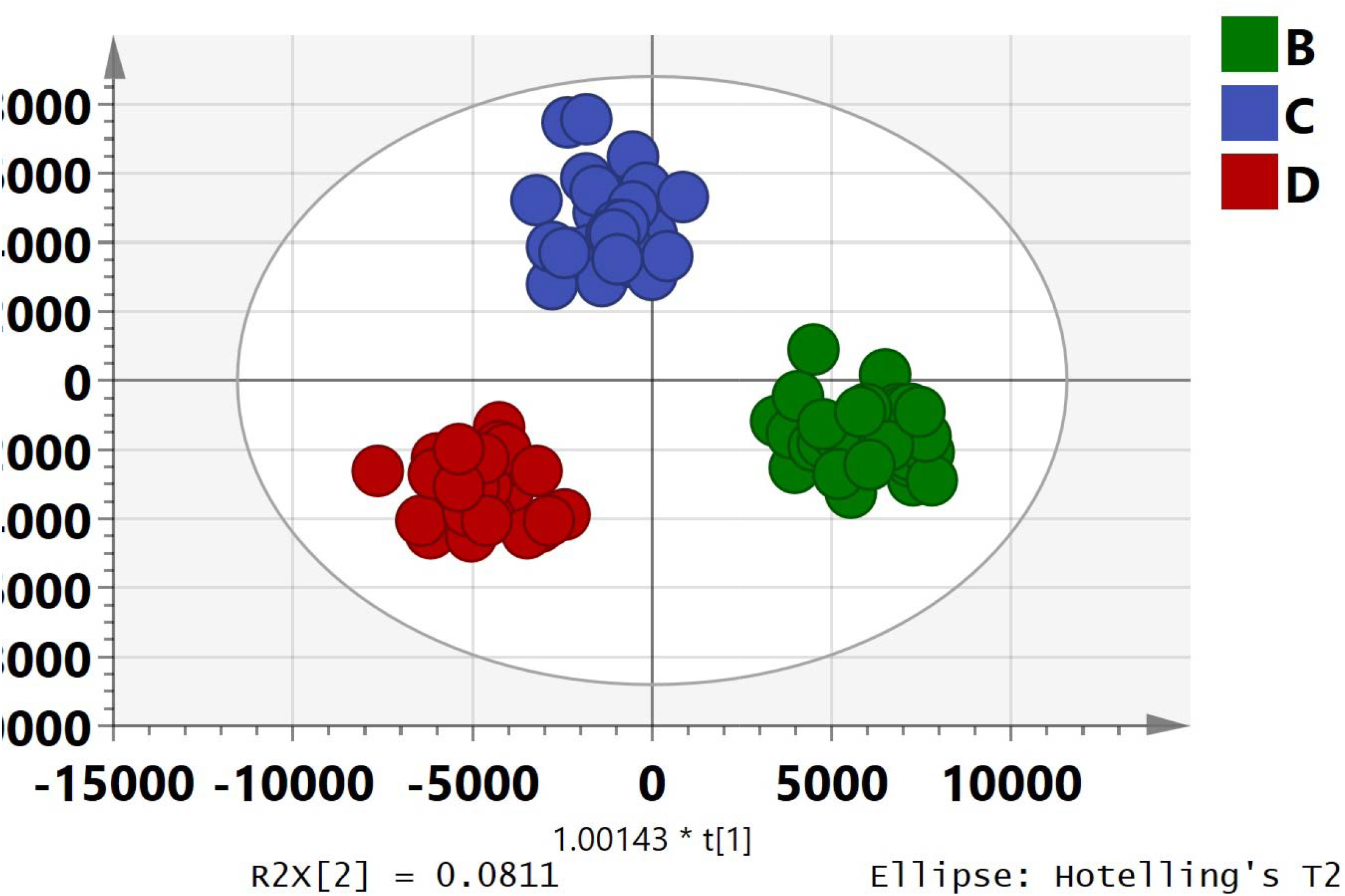
OPLS-DA plot for seasonal differentiation season B: May 2013; season C: October 2013; season D: March 2014

### Bioassays

Based on the cut-offs selected for each bioassay, extracts were classified having good, intermediate and poor activity. Table 1 gives the cut-offs and the number of extracts under these three categories for the individual bioassays.

The numerical data of bioassay was correlated with the HPLC data using suitable statistical analysis. In all assays except CT binding, use of PCA was unsuccessful for developing a model with statistical significance. Hence OPLS-DA or OPLS-RA was used with either all 90 extracts or a subset of data as discussed individually below. Table 2 summarizes the developed models for the individual bioassays.

**Table 2:** Summary of models developed for each bioassay showing statistical significance.

| Assay | Method | Number of extracts included |
| --- | --- | --- |
| **Successfully established models for** |  |  |
| **Antibacterial activity – *S. flexneri*** | OPLS-RA | 90 |
| **Antibacterial activity – *V. cholerae*** | OPLS-RA | 90 |
| **Invasion of EIEC** | OPLS-RA | 90 |
| **Cholera toxin (CT) production** | OPLS-RA | 90 |
| **CT binding** | OPLS-DA | 90 |
| **Models established after exclusions of confounders ( )** |  |  |
| **Adherence of EPEC** | OPLS-RA | 20 G + 18 B<br>(5 G + 1 B) |
| **Invasion of *S. flexneri*** | OPLS-RA | 20 G + 13 B<br>(8 G + 6 B) |
| **Labile toxin production** | OPLS-RA | 14 G + 23 B<br>(8 G + 12 B) |

OPLS-RA: Orthogonal Partial Least Square-Regression Analysis
OPLS-DA: Orthogonal Partial Least Square-Discriminant Analysis
G: Extract with Good activity; B: Extract with Bad/Poor activity

#### Antibacterial activity

Based on the cut-offs, 21 extracts exhibited good activity and 53 were associated with bad activity for antibacterial activity against *S. flexneri*. As no segregation was observed in the PCA plot with all 90 extracts, OPLS (regression) plot was further developed with all the extracts. As seen from Figure 4, this approach revealed significant segregation of good and poor activity extracts (*P* = 0.00851913; R^2^X= 0.538; Q2 (cum) = 0.217).

**Fig. 4:**
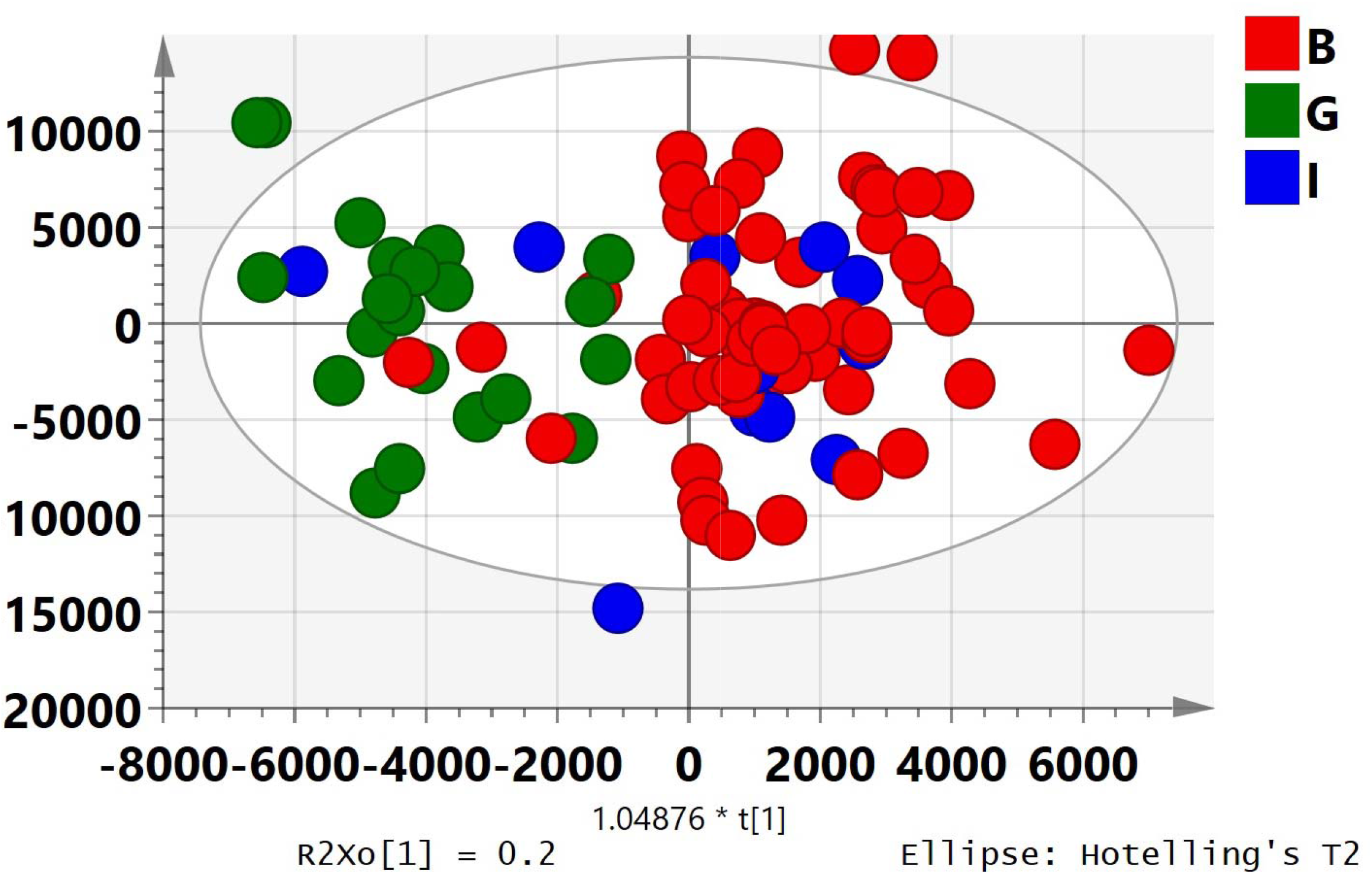
OPLS-Regression model for antibacterial activity against *Shigella flexneri* G: Good activity; B: Bad/Poor activity; I: Intermediate activity

With respect to antibacterial action against *V. cholerae,* though there were 45 extracts showing poor activity, only seven extracts displayed good activity. For correlating this biological activity with the HPLC data, similar to the antibacterial activity against *S. flexneri*, OPLS (regression analysis) plot was developed with all 90 extracts. Figure 5 shows significant segregation of good and poor activity extracts (*P* = 2.39×10^-6^; R^2^X= 0.66; Q2 (cum) = 0.449) based on this plot.

**Fig. 5:**
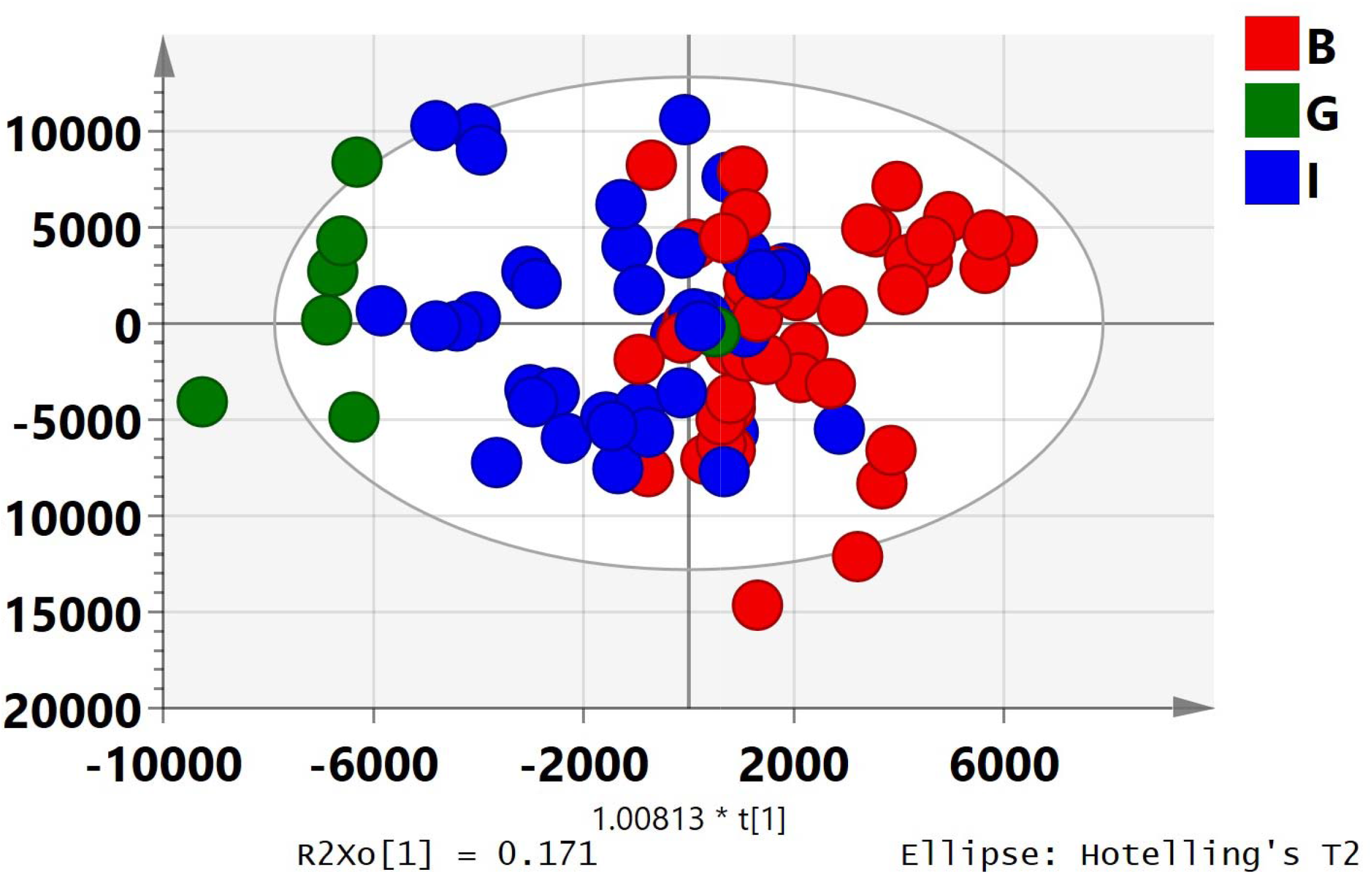
OPLS-Regression model for antibacterial activity against *Vibrio cholerae* G: Good activity; B: Bad/Poor activity; I: Intermediate activity

Though, antibacterial activity of the extracts was undertaken with *E. coli* strains (EPEC, ETEC) too, it was found that the two strains of bacteria were resistant to the extracts to a higher degree, showing only partial growth inhibition at a concentration of 1 mg/mL of the extract (data not shown). Hence, no further assays and any subsequent analysis were undertaken with *E. coli*.

#### Bacterial colonization

##### Adherence of EPEC

Of the 90 extracts, 25 and 19 extracts were found to have good and bad activity, respectively, for adherence of EPEC to HEp-2 cells. Unlike the antibacterial activities against *S. flexneri* and *V. cholerae*, the inclusion of all 90 samples did not develop a significant model using regression analysis (Figure 6a). Hence, further attempts to develop a significant model were made. An OPLS-RA model was then established based on the samples that depicted high and low inhibition. This model included 20 good activity extracts and 18 poor activity extracts (*P* = 0.0466, R^2^X= 0.325; Q2 (cum) = 0.248; Figure 6b). Thus, this model excluded five good and one poor activity extracts.

**Fig. 6a:**
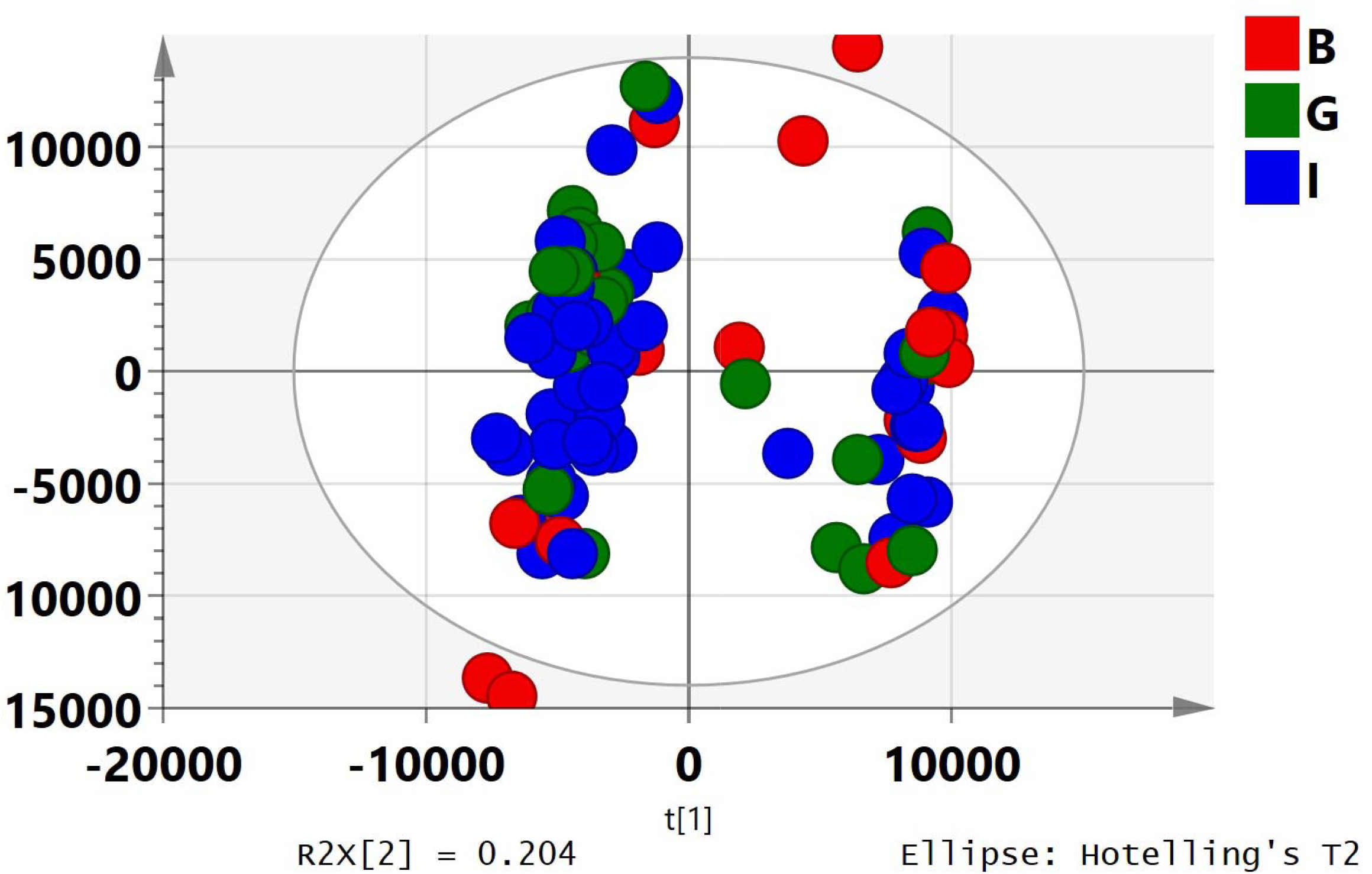
PCA model for effect on adherence of EPEC G: Good activity; B: Bad/Poor activity; I: Intermediate activity

**Fig. 6b:**
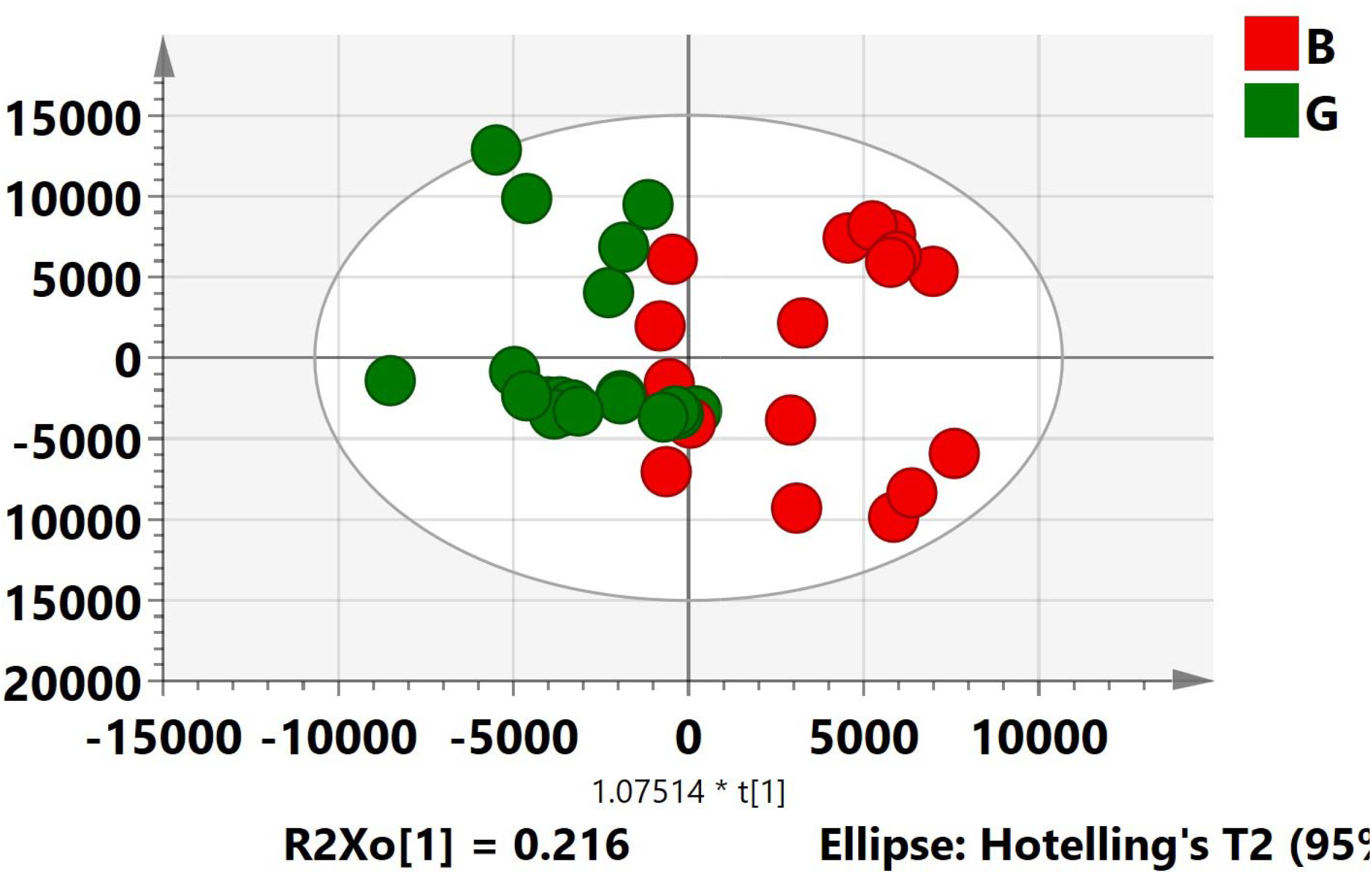
OPLS-Regression model developed for effect on adherence of EPEC G: Good activity; B: Bad/Poor activity

##### Invasion of EIEC

There were almost same numbers of extracts having good and poor activity for invasion of EIEC (22 and 21, respectively). For this activity, an OPLS-RA model (Figure 7) with the 90 extracts showed good segregation of good and poor activity extracts (*P* = 0.263; R^2^X= 0.12; Q2 (cum) = 107).

**Fig. 7:**
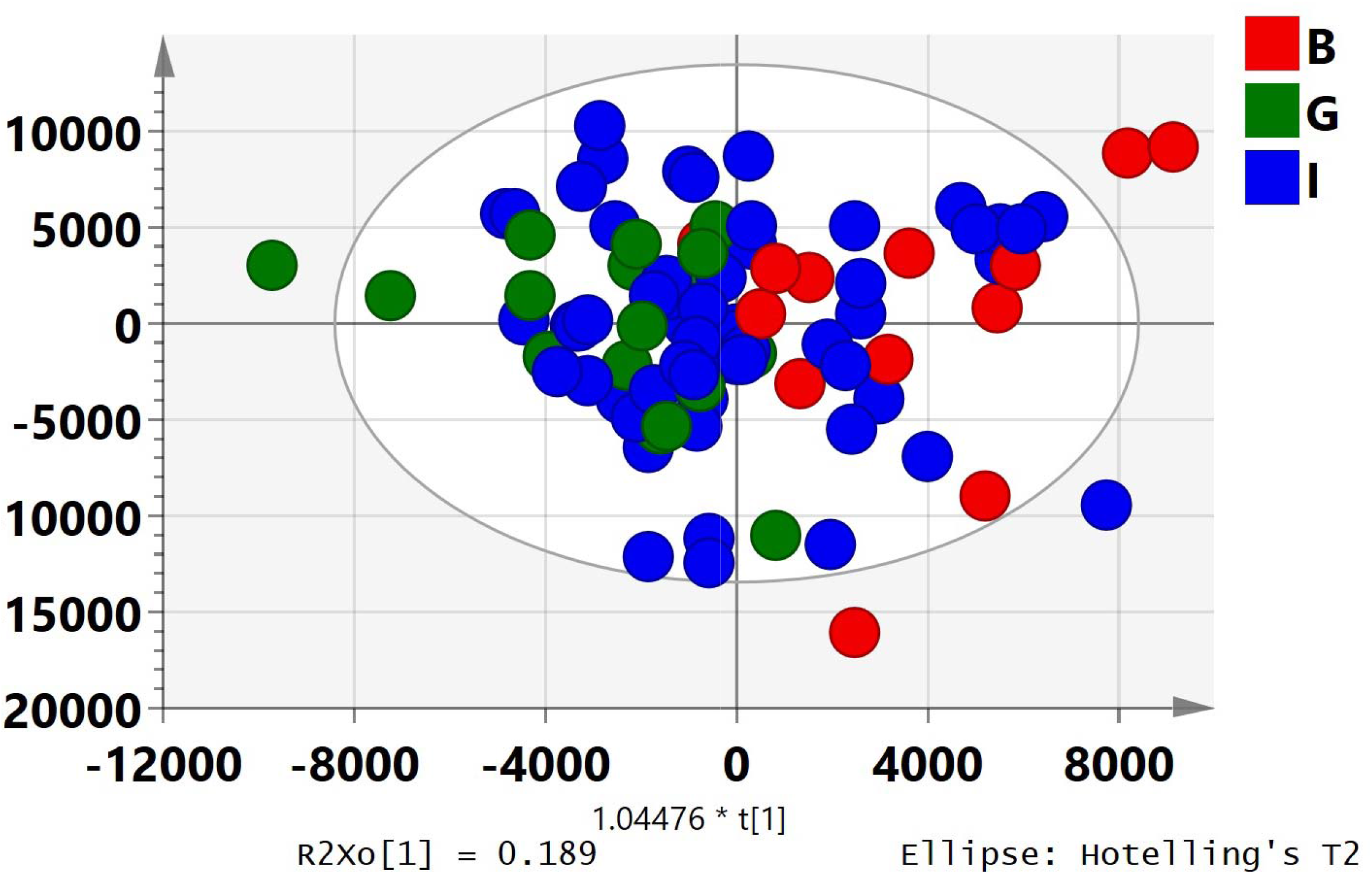
OPLS-Regression model for effect on invasion of EIEC G: Good activity; B: Bad/Poor activity; I: Intermediate activity

##### Invasion of *S. flexneri*

The number of extracts that could inhibit the invasion of *S. flexneri* to HEp-2 cells was 28; there were 19 extracts that showed poor inhibition of the bacterial invasion. For this assay, an OPLS-RA model with all 90 extracts did not yield any significance (Figure 8a). Hence, a sub set of 33 extracts (20 good and 13 poor activity extracts) after excluding confounders (8 good and 6 poor activity extracts) was used to develop a significant model (Figure 8b, *P* = 0.0114; R^2^X= 0.33; Q2 (cum)= 0.361).

**Fig. 8a:**
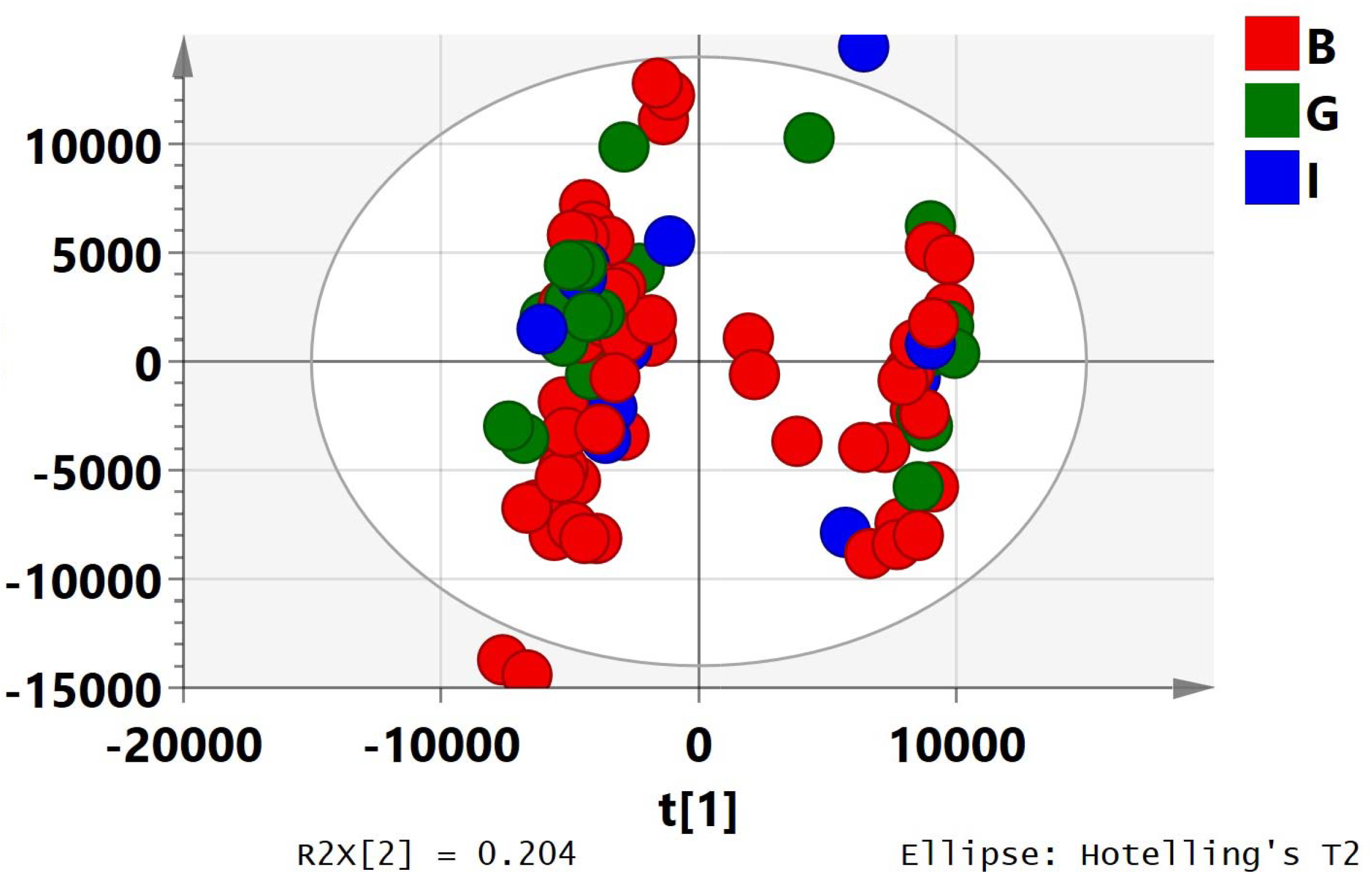
PCA model for effect on invasion of *Shigella flexneri* G: Good activity; B: Bad/Poor activity; I: Intermediate activity

**Fig. 8b:**
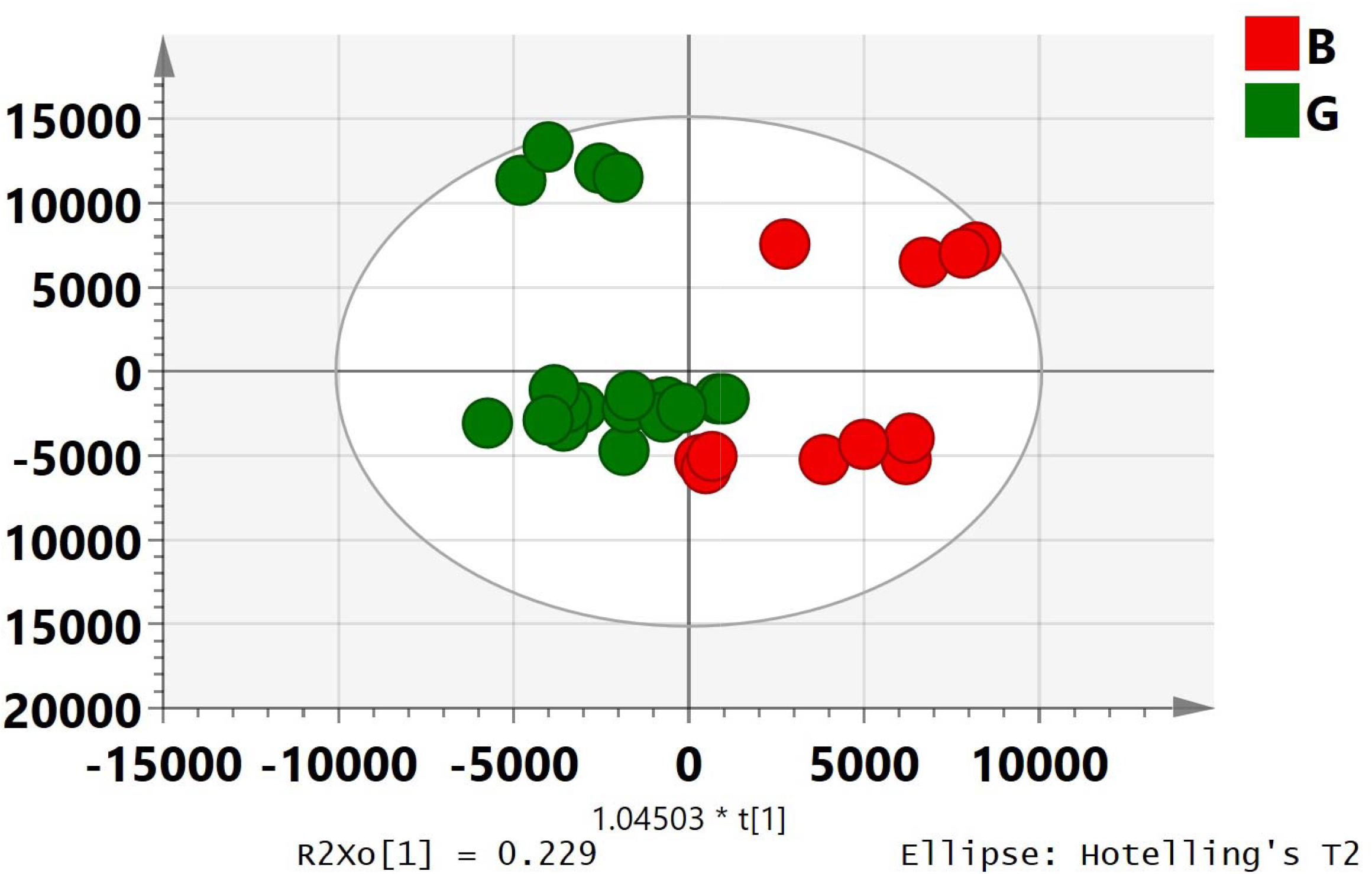
OPLS-Regression model developed for effect on invasion of *Shigella flexneri* G: Good activity; B: Bad/Poor activity

#### Bacterial enterotoxins

##### CT production

For this bioassay, 15 extracts showed good activity and 22 were noted to have poor activity. The OPLS-RA model revealed a good segregation of good and poor activity extracts (Figure 9, *P* = 0.00098; R^2^X= 0.286; Q2 (cum) = 0.194) using all 90 extracts.

**Fig. 9:**
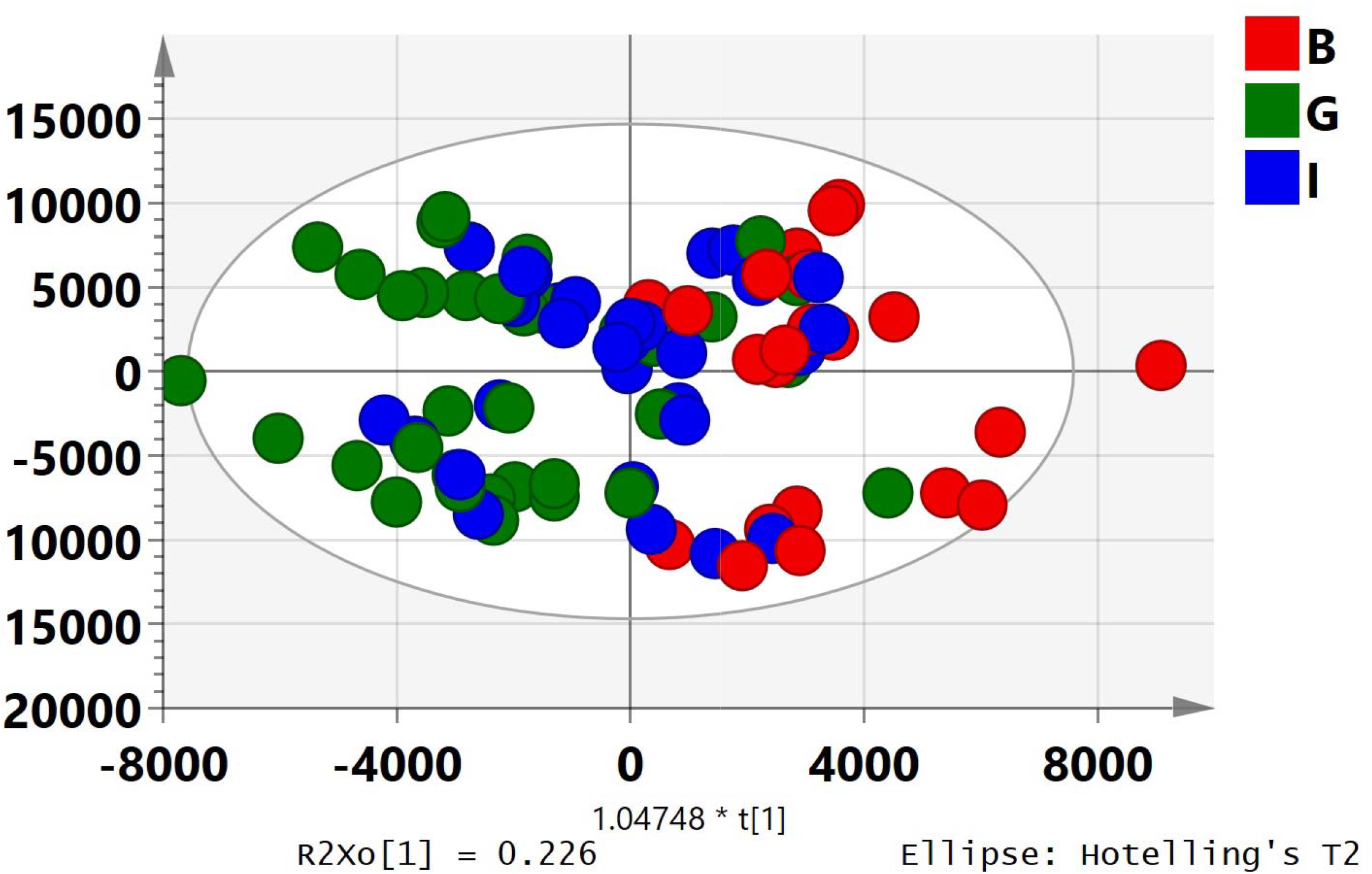
OPLS-Regression model for production of cholera toxin by *Vibrio cholerae* G: Good activity; B: Bad/Poor activity; I: Intermediate activity

##### CT binding

Of the 90 extracts, while 20 showed good inhibition, 26 could not inhibit the toxin. While correlating, this was the only assay wherein no significant model could be developed with PCA and OPLS (regression) using all 90 extracts. An OPLD-DA plot further developed, revealed a good segregation of good and poor activity extracts (*P* = 0.016; R^2^X= 0.297; Q2 (cum) = 0.111) with all 90 extracts as depicted in Figure 10.

**Fig. 10:**
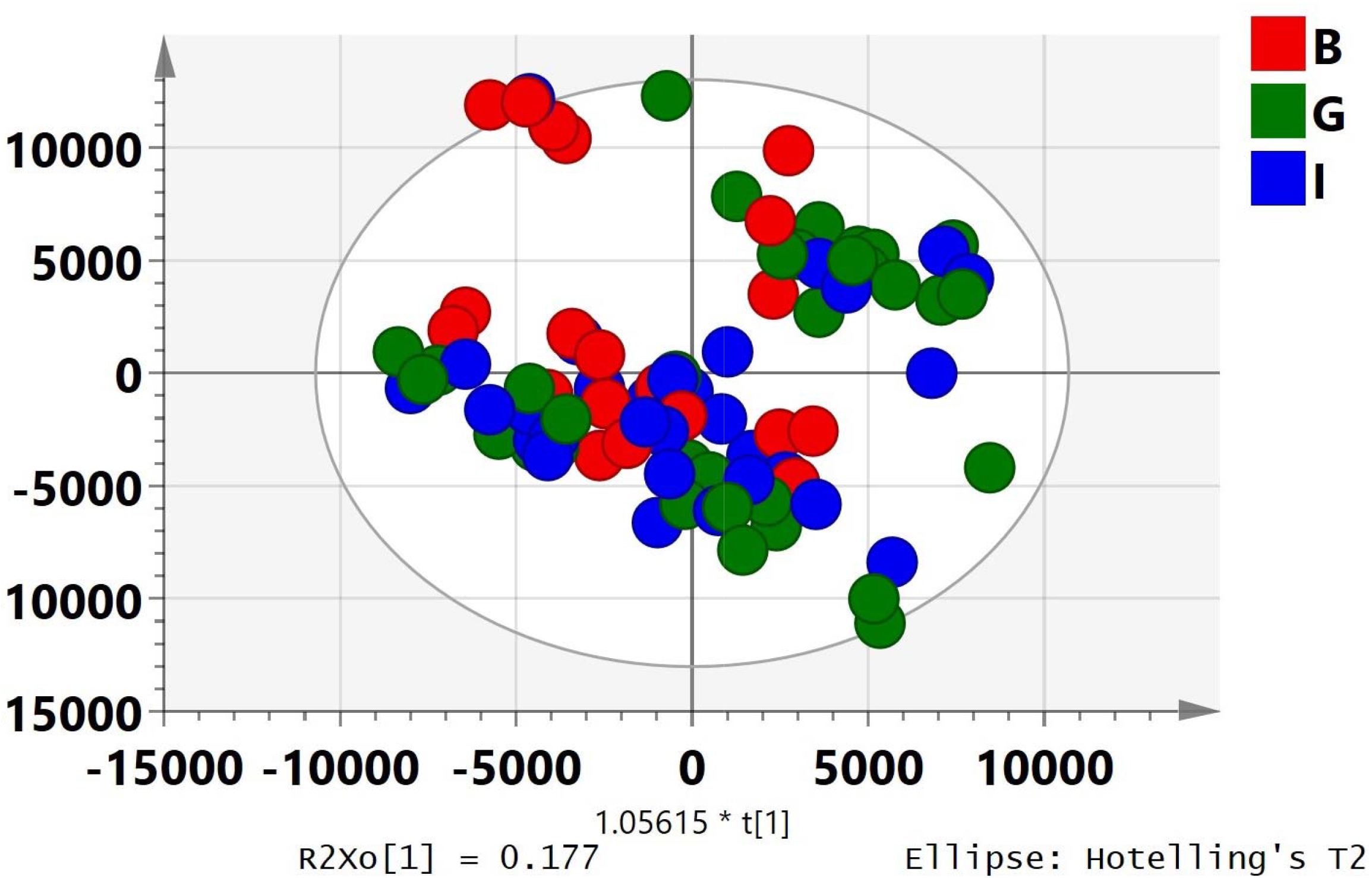
OPLS-DA model for binding of cholera toxin by *Vibrio cholerae* G: Good activity; B: Bad/Poor activity; I: Intermediate activity

##### LT production

Twenty-two extracts showed good inhibition of LT, and 35 had poor activity against this toxin. No significant model could be established using all 90 extracts (Figure 11a). Hence, an OPLS-RA model showing segregation of good and poor activity extracts (Figure 11b, *P* = 0.0356; R^2^X= 0.283; Q2 (cum) = 0.268) with 37 extracts (14 good and 23 poor activity extracts) following exclusion of 14 extracts (8 good and 12 poor) was successfully established.

**Fig. 11a:**
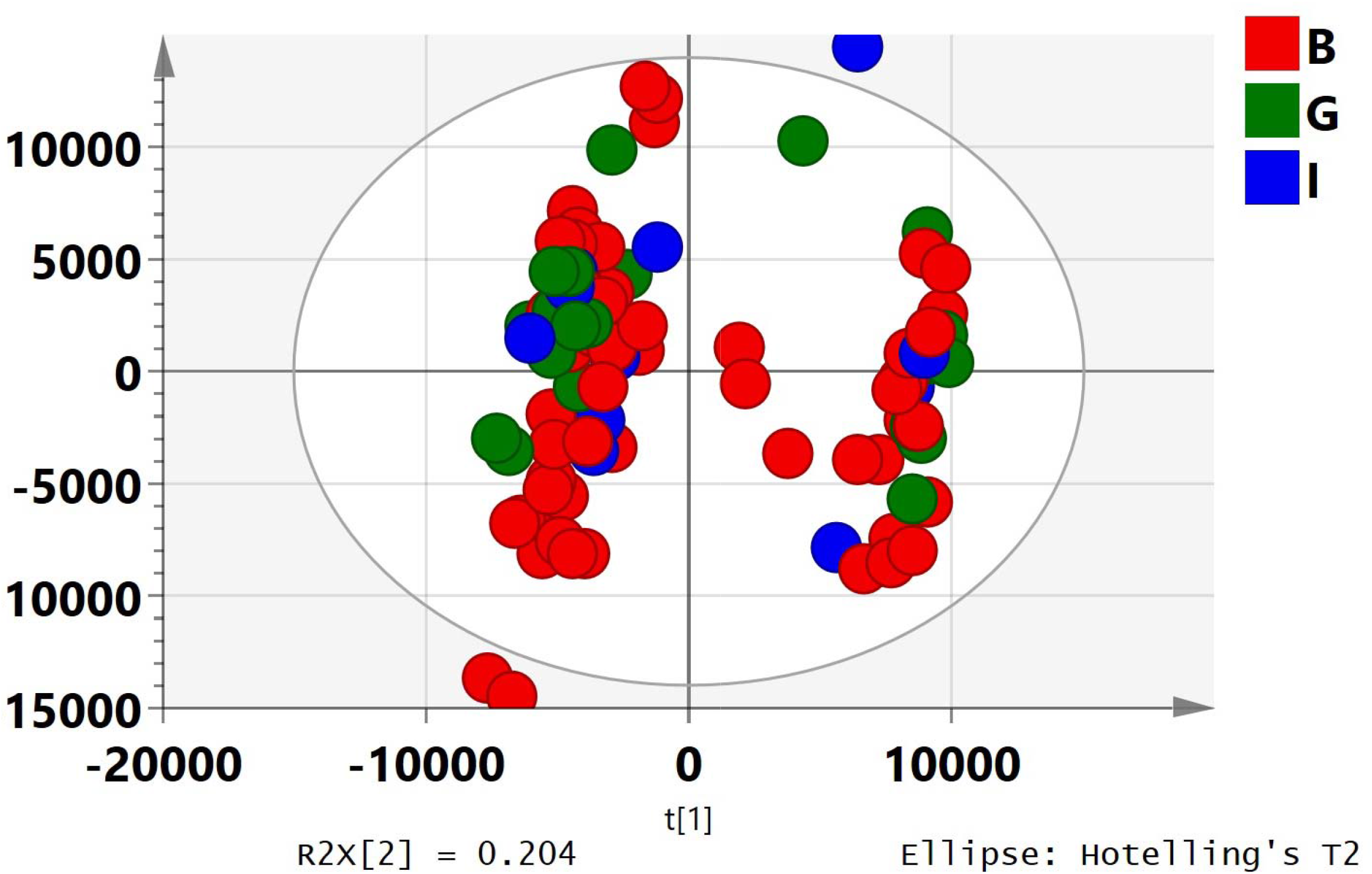
PCA model for effect on production of labile toxin G: Good activity; B: Bad/Poor activity; I: Intermediate activity

**Fig. 11b:**
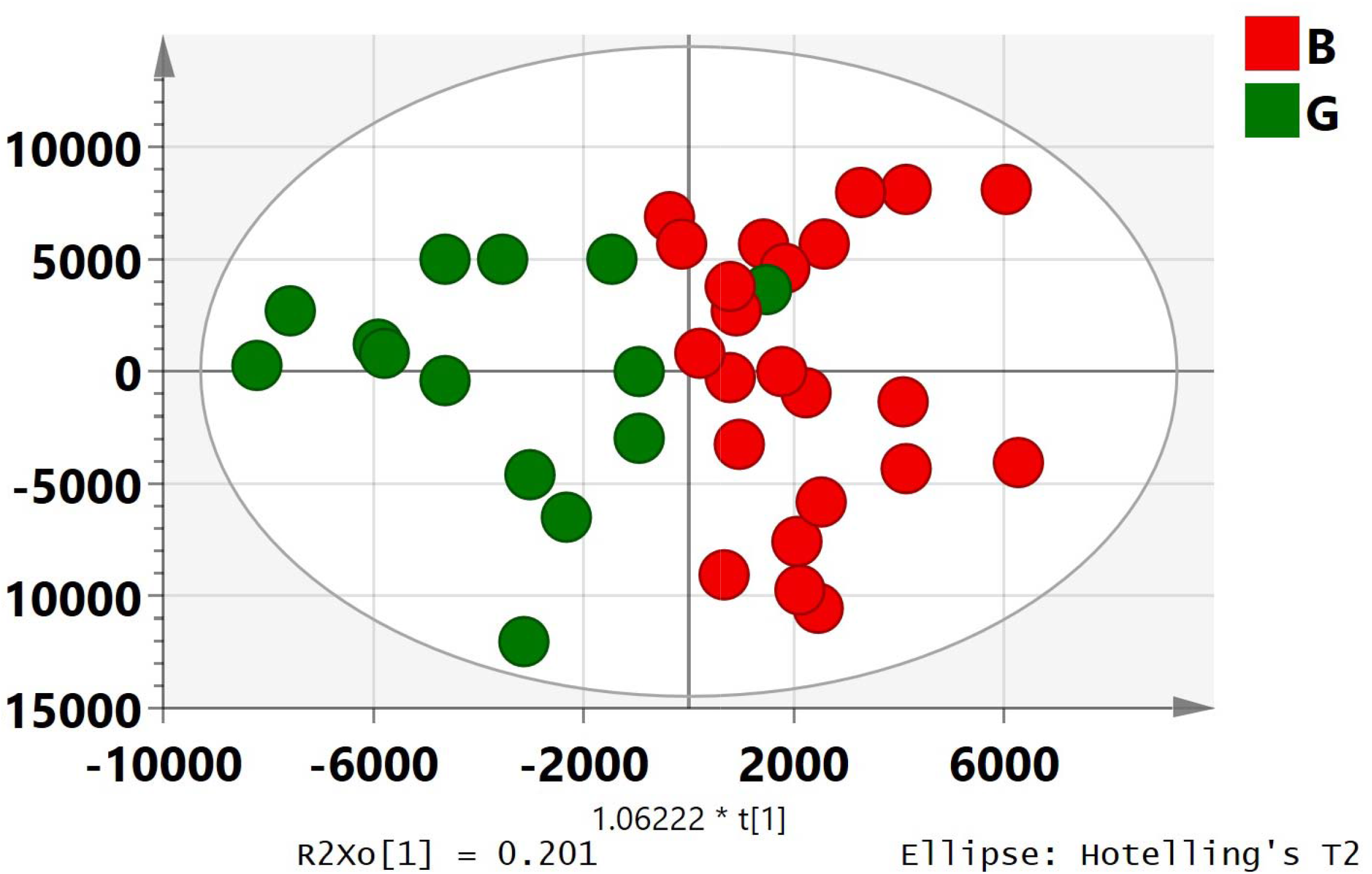
OPLS-Regression model developed for effect on production of labile toxin G: Good activity; B: Bad/Poor activity

## Discussion

An incredible increase in the global use of medicinal plants has led to the development of several concerns related to their standardization. Standardization of herbal medicine is a critical aspect to minimize batch to batch variability so as to ensure uniform efficacy and safety thereby supporting its acceptability. As a method of standardization of crude extracts, we have employed metabolomics using the antidiarrhoeal activity of guava leaves as an example.

Metabolomics is a promising and developing field of ‘omics’ research that is concerned with characterizing large numbers of metabolites. This field has emerged as an important tool in plant research. It has been documented to have a number of applications such as assessing quality, detecting adulterants, evaluating biological efficacy and determining optimum conditions for cultivation (Hall, 2006; Wolfender et al., 2013; Ning et al., 2013). Various spectroscopic, chromatographic or electrophoretic techniques such as NMR, Fourier Transformed IR (FTIR), HPLC, Mass Spectroscopy (MS), Liquid Chromatography-MS (LC-MS), Gas Chromatography-MS (GC-MS) have been used in metabolomic studies each having its advantages and disadvantages (Lajis et al, 2017).

In our previous study, we attempted to correlate ^1^H NMR spectra of guava leaf extract with the biological data of eight assays so as to develop a prototype for standardization. NMR spectroscopy was chosen as it is a reproducible method for metabolome analyses, involving simple sample preparation and lower measuring times (Ali et al. 2013). This spectroscopic method can be applied to complex mixtures of metabolites such as those seen in plants. However, a single peak in NMR may not correspond to a proton from a single molecule and hence identification of compounds is difficult. Moreover, the high end NMR equipment is available in only few laboratories/industries and its maintenance cost is expensive. On the other hand, though HPLC is not as sensitive and reproducible as the NMR technique, a single peak in HPLC chromatogram usually correlates to a single compound and thus can be used to obtain important information about individual constituents of crude extracts (Bertrand et al. 2016). HPLC equipment too is available in most of laboratories/industries and the maintenance cost is less. In the present study, we used HPLC coupled with metabolomics as a standardization approach.

Despite the plant material being collected from open field conditions and a high degree of variability expected, the metabolomics approach used in the present study could successfully correlate the HPLC chromatograms to most of the bioassays employed. Thus, HPLC-based metabolomics does seem a promising approach as it could help differentiate guava extracts not only on the basis of the regional and seasonal collection but also on the bio-assays carried out. It would be interesting to further analyze and evaluate the data for understanding the compound(s) present in the guava extract that could be used as indicators of the anti-diarrhoeal activity of this promising plant and further used for standardization of the extract.

## Funding

This work was supported by Zoetis Pharmaceuticals Research Private Limited, India.

## Acknowledgments

The authors thank Dr. P. Tetali for collection the guava leaves used in the study. The authors are also grateful to Dr. Purnendu Roy-Choudhury (TCG Life Sciences, Kolkata) for facilitating the acquisition of HPLC fingerprinting of the extracts. The efforts of Dr. Ajit Datar and Mr. Shailesh Damale (Shimadzu, Mumbai) towards conversion of acquired HPLC data to numeric format are also acknowledged.

## Notes

### Competing Interest Statement

The authors have declared no competing interest.

### Summary of Updates

The figures were not numbered in the previous submission. In this revision only the figures have been assigned numbers.

